# Dispersal dynamics and local filtering vary with climate across a grassland landscape

**DOI:** 10.1101/567586

**Authors:** John Guittar, Deborah Goldberg, Kari Klanderud, Astrid Berge, Marta Ramírez Boixaderas, Eric Meineri, Joachim Töpper, Vigdis Vandvik

## Abstract

Dispersal dynamics and local filtering interactively generate regional vegetation patterns, but empirical evidence of their combined influence in nature is scarce, representing a missing link between our theoretical understanding of community assembly and real-world observation. Here, we compare seed and adult plant communities at twelve grassland sites with different climates in southern Norway to explore the degree to which community membership is shaped by dispersal limitation and local niche-based filtering, and how this varies with climate. To do this, we first divide species at each site into two groups: “locally-transient” species, which occur as seeds but are rare or absent as adults (i.e., they arrive but are filtered out), or “locally-persistent” species, which occur consistently as adults in annual vegetation surveys. We then ask questions to reveal where, when, why, and how locally-transient species are systematically disfavored during community assembly. Our results led to four main conclusions: (1) the strength of local filtering on community membership increased with temperature, (2) surprisingly, local filtering was stronger during seedling emergence than during seedling establishment, (3) climate-based niche differences drove differential performance among species, especially for seeds dispersing outside of their realized climate niches into more stressful (colder and drier) climates, and (4) locally-transient species had traits that may made them better dispersers (smaller seeds) but poorer competitors for light (shorter statures, less persistent clonal connections) than locally-persistent species, providing a potential explanation for why they arrived to new sites but failed to establish persistent adult populations. Our study is one of the first to combine seed, seedling, and adult survey data across multiple sites with different climates to provide a rigorous empirical evaluation of the combined influence of dispersal limitation and local filtering on the generation and maintenance of climate-associated vegetation patterns.

## INTRODUCTION

Community assembly theory in a landscape or metacommunity context assumes an important role for regional dispersal dynamics, either alone (i.e., neutral dynamics) or in conjunction with niche-based performance differences (Leibold et al. 2004, Alexander et al. 2012). While empirical evidence has demonstrated that dispersal and niche-based differences can each influence community membership on their own (Choler et al. 2001, Ehrlén et al. 2006, Armas et al. 2011, Laliberté et al. 2014), there is a lack of data detailing how these processes interact in nature, and how their combined influence changes across environments, representing a missing link between theoretical assumptions and real-world observations. This lack of empirical data raises potential problems when trying to model community responses to disturbances like climate change, because different species may have different dispersal abilities and/or different environmental constraints to expansion, leading to variable responses (Graae et al. 2017).

The lack of regional dispersal data in community assembly research is particularly acute, and this is due in large part to logistical challenges. Plant propagules are often tiny, numerous, difficult to identify, capable of traveling great distances, and can remain dormant in the soil for years prior to germination (Baskin and Baskin 1998, Vandvik et al. 2016). Some researchers have sidestepped these difficulties by inferring dispersal patterns using indirect methods (Alexander et al. 2012). The “nearest-neighbor” approach, for example, assumes connectivity in a metacommunity to be proportional to inter-patch distance (Calabrese and Fagan 2004, Jacobson and Peres-Neto 2010). Other approaches infer dispersal limitation from historical population shifts over periods of climate change (Kelly and Goulden 2008, Bertrand et al. 2011) or by analyzing population genetic structure. These indirect methods are useful when modeling species distributions from a phenomenological standpoint, but fall far short of evaluating whether particular species arrive or not to a given site (Calabrese and Fagan 2004). The small number of studies that have directly assessed dispersal patterns, either by manually marking seeds (e.g., Xiao et al. 2006) or connecting seeds to parents using parentage analyses (e.g., Cain et al. 2000), are often conducted for single species and/or over short distances, and are of limited use when considering potential responses of dozens of species at landscape scales (Nathan and Muller-Landau 2000).

Here, we compare compositions of seed rain, seed bank, seedling, and adult plant communities at sites along temperature and precipitation gradients to shed light on how seed dispersal and local filtering interact *in situ* to shape community membership and maintain regional vegetation a patterns. First, to evaluate the strength of local filtering processes, we divide the species at each site into two groups: “locally-transient” species, which occur as seeds but are rare or absent as adults, and “locally-persistent” species, which occur consistently as adults in annual vegetation surveys. Unlike the distinction between “core” and “satellite” species *sensu* Hanski (1982), our framework allows species status to vary by site, rather than be defined regionally. Next, we ask questions to elucidate where, when, why, and how locally-transient species are selectively disfavored during community assembly, and how this process varies by climate. We also use our adult vegetation surveys to infer which locally-transient species have dispersed outside of their realized climate niches (i.e., are outside of the range of climates where we know they persist as adults), and how their realized climate niches compare to local climate conditions (i.e., if they have dispersed into warmer/wetter/cooler/drier climates). If we observe strong performance differences between locally-transient and locally-persistent species at a site, then we can conclude that local filtering processes (in combination with dispersal processes) play an important role in governing community membership at that site. If we do not observe strong performance differences, then we can conclude that dispersal dynamics (and not local filtering processes) primarily govern community membership.

Our study takes place in a network of 12 alpine and subalpine grassland sites in southern Norway, a region with unusually high spatial climate variability. The sites were selected according to their mean summer temperatures and mean annual rainfalls such that they form an orthogonal climate grid, facilitating independent assessment of these two important climate drivers in community assembly. Prior work used subsets of the data used in this study to compare diversity patterns in the seed bank and mature vegetation (Vandvik et al. 2016), to understand how trait-based community composition varies with climate (Guittar et al. 2016), and to evaluate the relative balance of competition and facilitation in seedling recruitment (Klanderud et al. 2017). We combine these previously published data with new data on seed rain and seedling survival to ask how regional dispersal dynamics and local filtering interactively govern community membership and regional vegetation patterns. After dividing species into locally-transient and locally-persistent groups at each site, we ask the following specific questions:

1. To what degree is community membership shaped by local filtering (i.e., what fraction of the seeds at a site are of locally-transient species), and how does the strength of local filtering vary with climate?
2. Are locally-transient species disfavored because they fail to emerge as seedlings, fail to establish, fail to compete as adults, or a combination of the three?
3. Are locally-transient species disfavored because they dispersed outside of their realized climate niches, indicating a role for climate-based niche filtering?
4. Are locally-transient species disfavored because they differ systematically from locally-persistent species in their functional traits, offering mechanistic hypotheses for their selective removal?

We conclude with a discussion of what our results mean for how these mountain grasslands will likely respond to climate change.

## MATERIALS AND METHODS

### Study area

The study area comprises 12 semi-natural calcareous grassland sites in southern Norway that host at least 144 non-woody vascular plant species at the adult life stage, and at least 126 at the seed stage (Appendix S1: Table S1). Sites have similar bedrock, land use histories, slopes of approximately 20°, and southwest aspects, but differ in mean summer temperature, defined as the mean temperature of the four warmest months per year, and/or mean annual precipitation, such that they form a grid with approximately orthogonal climate axes (Fig. 1). Interannual variation in mean summer temperature and annual precipitation at each site over the sampling period (2008 – 2013) was modest, with standard deviations of 0.79 °C and 238 mm, respectively, on average across sites. Each site has five blocks, placed in representative areas of grassland vegetation within a 75 – 200 m^2^ area that varies due to local topography and accessibility. Each block has a variable number of 25 x 25 cm plots used for the data sets described below, as well as other experiments and surveys. Blocks were protected from grazers with electric fences and manually mowed once a year to evenly simulate biomass loss due to grazing. Taxonomic identifications follow Lid & Lid (2007). Woody species and bryophytes were excluded from all analyses.

**Figure 1.**
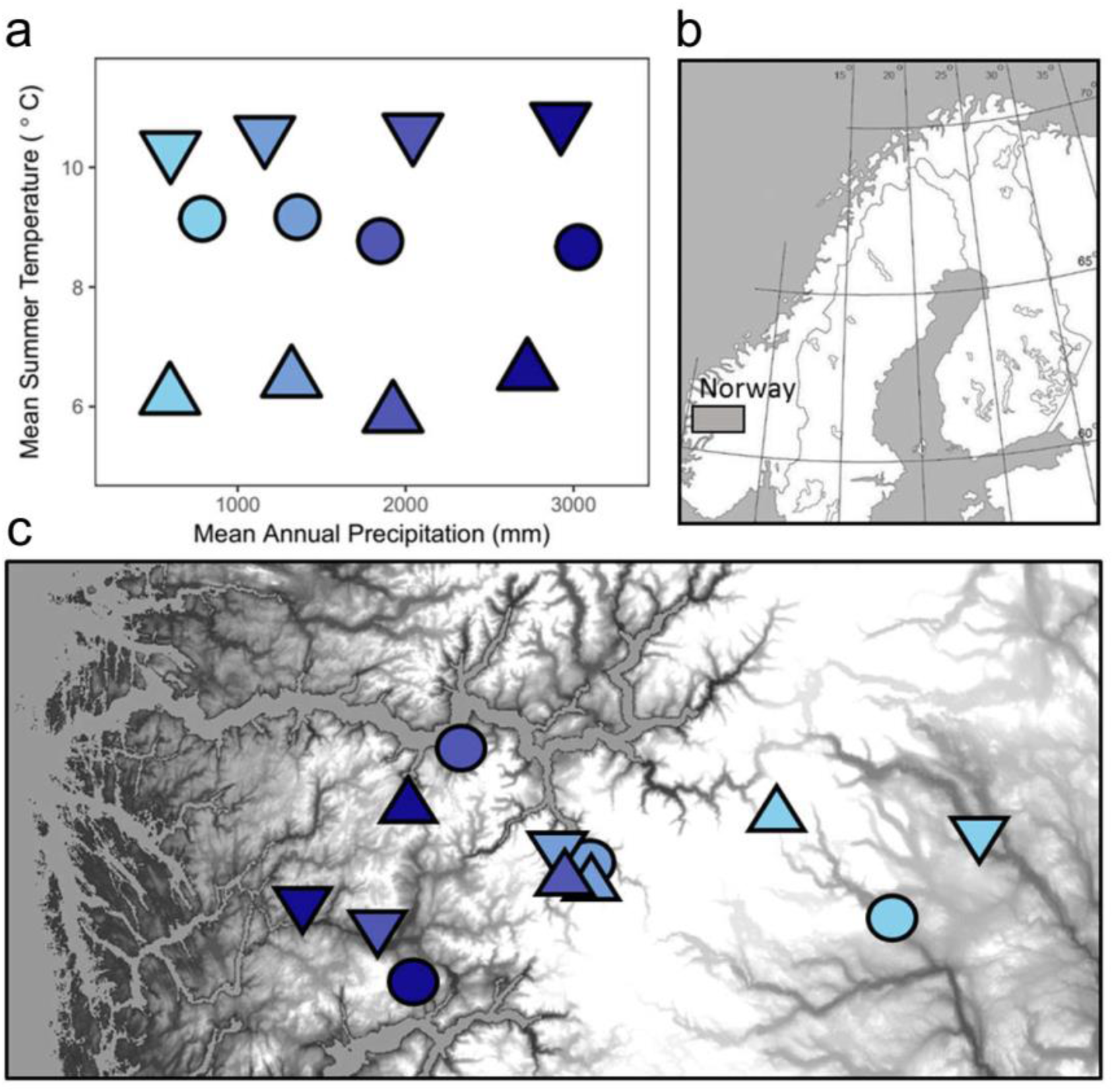
Climates (a) and locations (b and c) of the twelve field sites in southern Norway. Mean summer temperature is defined as the warmest four months at each site. Grayscale shading in c reflects altitude, which covaries closely with mean summer temperature.

### Seed rain data

We collected seed rain over two periods to target winter (September 2009 to June 2010) and summer (June 2010 to September 2010) seed deposition. We trapped seeds in 25 x 25 cm artificial turfs placed in gaps about 50 cm from seedling monitoring gaps (see below) in four blocks at each site, for a total of 48 traps across all sites. The small synthetic filaments in artificial turfs effectively catch and retain small particles like seeds. Turfs were gathered and flushed with water to free collected seeds. The rinse water was passed through 500 µm and 125 µm diameter sieves to discriminate seeds by size and remove debris. Seeds were counted and identified taxonomically using a stereomicroscope. We included fruits, bulbils, and viviparous seeds in our working definition of “seeds.” Rates of seed predation in artificial turfs are likely similar to natural rates experienced by seeds on the natural soil surface, thus seed predation should not bias our results.

### Seed bank data

To characterize seed bank diversity we haphazardly selected one 64 x 64 cm plot at each site and excavated soil to a depth of 3 cm in October 2008. Because the seed bank survey area at each site (0.41 m^2^) was larger than the total areas used for seed rain and seedling surveys at each site (four 25 x 25 cm plots, 0.25 m^2^), we randomly subsampled 61 % (0.25 m^2^ / 0.41 m^2^) of the original seed bank community *in silico* and discarded the remainder to standardize sampling effort across data types at each site. To exclude the seeds from that year’s seed rain from our seed bank surveys, we removed aboveground vegetation, including moss and litter. Soil samples were stored for three months at 2 – 4 °C and ambient moisture. Soil samples were then sowed into a standard mixture of sterile subsoil and placed in 30 x 60 cm trays. The trays were incubated in a greenhouse with a diurnal cycle of 16 hours light (25 °C) and 8 hours darkness (15 °C). The diurnal cycle was continued for four months, followed by six months of cold stratification in darkness (4 °C), followed by another four-month period of diurnal cycling. Emerging seedlings were counted and removed once identifiable to species. This method of characterizing the seed bank community effectively accounts for seed viability because non-viable seeds would not have emerged as seedlings.

### Seedling data

Like most perennial grasslands, seedling recruitment in our system is highly dependent on disturbances and occurs only rarely in intact vegetation due to strong competitive effects from adult plants (Silvertown and Smith 1989, Eriksson 1989, Bullock et al. 1995, Vandvik 2004, Berge 2010, Klanderud et al. 2017). We therefore monitored seedlings in experimental gaps where they were relatively free from competitive effects of adult plants, but still exposed to environmental stress and other biotic interactions, such as herbivory, disease, and potential resource competition among seedlings. One 25 x 25 cm gap was created in each of four blocks at each site in spring 2009, for a total of four gaps per site and 48 gaps overall. The gaps were made by cutting along the inner edges of a metal frame mounted in metal pipes marking the corners of the plot, and peeling away the natural vegetation and its thickly interwoven root mat. Seeds and topsoil were returned to gaps by vigorously shaking excavated vegetation and passing it through a 4 mm sieve to remove plant remains. Emerged seedlings in the plots were ID-tagged in one of three censuses (late summer 2009, early summer 2010, late summer 2010) using numbered plastic toothpicks and plot coordinates. About 70 % of seedlings were identifiable to species; the remaining 30 % were unidentifiable or died before they could be identified and were lumped into two generic groups for graminoids and forbs. Seedlings were differentiated from emergent clonal ramets by looking for cotyledons or signs of above- or below- ground connections. Seedling survival and establishment were recorded twice yearly from spring 2010 to spring 2012. We approximated seedling emergence rates by dividing the density of emerged seedlings by the sum of seed rain and seed bank densities. We marked seedlings established when they had grown to a size greater than what could be derived exclusively from their maternal subsidies, which we estimated to be when stems were longer than 2 cm, and if a forb, also when they had grown their first non-cotyledonous leaves. Although we took pains to census each site at times of peak seedling emergence, some seedlings may have emerged, died, and disappeared before ever being recorded; even if this where the case, however, we contend that it is unlikely to influence our overall conclusions because the factors discouraging seedling survival during the earliest stages of establishment are likely the same as those discouraging seedling emergence (i.e., late spring frosts), and are distinct from the factors influencing seedling survival establishment (i.e., drought, competition for light, predation, early fall frosts, and late spring frosts in the year after seedling emergence).

### Mature vegetation data

We surveyed mature vegetation at peak biomass (July and August) in 2009, 2011, 2012, and 2013. At each site, we used two 25 x 25 cm plots in each of five blocks at each site, for a total of 10 plots per site (except for the third-wettest second-warmest site, which only had nine plots), and 119 plots overall. These plots were controls for a transplant experiment and included five undisturbed controls and five transplant controls, i.e., turfs dug up and replaced in the same location. The two types of controls did not differ in species composition or any other aspect of community structure at any of the survey periods (Guittar et al. 2016). We visually estimated the percent cover of individual species in each plot using a 5 x 5 cm grid overlay, and then pooled the data by site. To ensure that site vegetation was not undergoing successional changes that could bias our conclusions, we used an NMDS ordination to confirm that site composition did not change substantively between years (Appendix S1: Fig. S1). Stage-specific species abundance data are provided as a supplementary file.

### Trait data

We used four commonly used plant traits, and four traits related to clonal growth strategy, each with hypothesized associations to dispersal ability and/or resource competition. Seed mass (mg), a reflection of species regeneration strategy (Kraft et al. 2008, Cornwell and Ackerly 2009), was drawn from the Seed Information Database (Royal Botanic Gardens Kew 2014). Maximum canopy height (m) data, which relates to both dispersal ability and light competition (Westoby 1998, Falster and Westoby 2003), were mined from Lid and Lid (2007). Leaf area (mm^2^) and specific leaf area (SLA; m^2^/kg), two traits indicative of where species fall along a continuum of slow-to-fast resource use strategies (Reich et al. 1997, Ackerly and Reich 1999, Reich 2014), were estimated using a combination of field data (Guittar et al. 2016) and data from the LEDA online trait database (Kleyer et al. 2008). Leaf area, SLA, maximum height, and seed mass values were log-transformed. Clonal traits included the number of offspring per parent per year (“0” = 1 offspring; “1” = ≥ 2 offspring), persistence of plant–offspring connections (“0” = < 2 years; “1” = ≥ 2 years), rate of lateral spread (“0” = ≤ 1 cm/year; “1” = > 1 cm/year), and number of buds per ramet (an integer score ranging from “1” = few buds either belowground or aboveground, to “8” = many buds both below and aboveground). Clonal attributes are thought to help plants integrate over spatially heterogeneous resources (Eilts et al. 2011), recover from disturbances (Klimešová and Klimeš 2007), and provide sustained maternal subsidies to new ramets as they grow horizontally and vie for local establishment (Herben and Wildová 2012). Clonal trait data were drawn from Klimeš and Klimešová (1999) and converted from categorical to quantitative formats to enable calculations of community means. Trait data are provided as a supplementary file.

### Assigning local species status

Each species observed at each site at any life stage was labeled as “locally-persistent” if adults were recorded in more than half (i.e., at least three of four) of the site vegetation surveys conducted from 2009 to 2013, or otherwise labeled “locally-transient.” In using this cutoff, we aim to understand the mechanisms leading a species to be a consistent member of a given community, not whether it can occur there at all. That is, an individual of a locally-transient species need not be a first generation immigrant, but could be a second-or-more generation immigrant, so long as it has evidently unable to establish a locally-persistent adult population. Because locally-transient/locally-persistent species status assignments were potentially sensitive to the depth at which we characterized local site community composition, we used rarefaction to verify that we had sufficiently surveyed the mature vegetation such that the number of locally-persistent species observed at each site had stabilized (Appendix S1: Fig. S2). In addition, we re-ran our analysis with all possible locally-transient/locally-persistent cutoffs to assess the sensitivity of our conclusions to our methodology.

### Assigning putative climate origins

For each locally-transient species at each site, we identified the sites and climates where we knew it had persistent adult populations, and then used these to infer whether it was dispersing outside of its realized climate niche. If a locally-transient species at a given site had persistent adult populations at other sites with similar temperatures or precipitations, we assumed it dispersed from a site with the “same temperature” or the “same precipitation,” such as from a neighboring site, e.g.., one with similar climate but with potentially different topographical, edaphic, or biotic characteristics. If a locally-transient species at a given site had persistent adult populations only at sites with warmer/cooler or wetter/drier conditions, we then assumed it dispersed from a warmer/cooler or wetter/drier site (i.e., it dispersed *into* a cooler/warmer or drier/wetter site). Again, we are not assuming that a given seed of a locally-transient species literally dispersed from a given climate, but that the seed is occurring outside of the climate range at which it is a common and persistent community member, and is presumably on the edge of being locally excluded.

### Statistical approach

We used linear regressions to test for baseline trends in the total abundance and richness of locally-transient and locally-persistent species in the seed rain, the seed bank, and the two combined along temperature and precipitation gradients. Unlike other data in this study, seed bank data were not collected in replicate across blocks within a site, thus our analyses were necessarily performed at a site-level resolution (N = 12). Each linear regression model assumes normally distributed errors and takes the form of *y_j_* ∼ MAP*_j_* or *y_j_* ∼ + MST*_j,_* where *y_j_* is the response variable being examined at site *j*, and MAP*_j_* and MST*_j_* are the mean annual precipitation (centered to zero) and mean summer temperature (centered to zero) at site *j*. We also used linear regressions to test for baseline trends in species richness in the mature vegetation with temperature and precipitation.

We used four sets of generalized linear models (GLMs) to test predictors of differential species performance during seedling emergence and seedling establishment. The dependent variable for the seedling emergence GLMs was the number of emerged seedlings of each species at each site (N =), and the dependent variable for the seedling establishment GLMs was the number of established seedlings of each species at each site. The GLMs used negative binomial error distributions and log link functions. Because each site has a unique combination of temperature and precipitation values (i.e., there is no nestedness), it was not appropriate to include site membership as a random effect while also testing for the effects of climate. When using seed and seedling numbers as predictors of emergence and establishment, respectively, we normalized their highly skewed abundance distributions with Yeo-Johnson transformations, which are similar to Box-Cox transformations but can be used with zeros. The alpha values used for the Yeo-Johnson transformations were those which yielded the most normal distributions, as quantified by Shapiro-Wilk normality tests; specifically, the alpha used to transform seed abundance data was -0.204 and the alpha used to transform emerged seedling data was -0.35.

In the first set of GLMs (“null” models), local species abundances in the combined seed rain and seed bank are used to predict local species abundances of emerged seedlings, and local species abundances of emerged seedlings predict local species abundances of established seedlings. In other words, our null expectation was that all seeds were equally likely to emerge, and all seedlings were equally likely to establish. In the second set of GLMs (“site climate” models), we added model terms for local site mean summer temperature and mean annual precipitation to evaluate how these climate variables affected seedling emergence and seedling establishment rates.

In the third set of GLMs (“site climate + species status” models), we evaluated how well locally-transient/locally-persistent species status predicted performance differences during seedling emergence and seedling establishment. Specifically, we added model terms specifying the status of each species at each site, and interactions between species status and local climate. We formally modeled the number of emerged seedlings *g* for species *i* at site *j* as

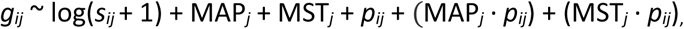

where *s_ij_* is the abundance of seeds (seed rain + seed bank) of species *i* at site *j*, *p_ij_* is a factor indicating species status, and MAP*_j_* and MST*_j_* are as described above. Likewise, we modeled the number of established seedlings *e* for species *i* at site *j* as

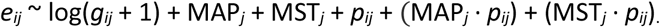

In the fourth and final set of GLMs (“site climate + species status + species climate origin” models), we dropped the model term for locally-transient/locally-persistent species status, *p_ij_*, and replaced it with a new term, *o_ij_*, denoting the putative temperature/precipitation origins of each locally-transient species *i* at site *j*. Specifically, the *o_ij_* term was used to tag each species at each site with one of five labels: (1) locally-persistent, (2) locally-transient but likely dispersed from an adjacent site with similar temperature/precipitation, (3) locally-transient and likely dispersed from a cooler/drier site (i.e., into a warmer/wetter site), (4) locally-transient and likely dispersed from a warmer/wetter site (i.e., into a cooler/drier site), (5) locally-transient with no locally-persistent populations at any of our grassland sites (i.e., an ‘unknown’ climate preference). We dropped the interaction terms from these GLMs to avoid excessive model complexity. Formally, we modelled the number of emerged seedlings *g* for species *i* at site *j* as

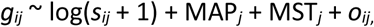

and the number of established seedlings *e* for species *i* at site *j* as

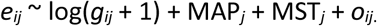

Finally, we asked if systematic differences in the traits of locally-transient and locally-persistent species at each site offered mechanistic explanations for performance differences between groups. To do this, we averaged the trait values of all species (not weighted by their relative abundances) in the combined seed rain and seed bank at each site, grouped by local-transient/locally-persistent species status. We used paired t-tests, paired by site, to identify which traits, if any, differed consistently between locally-transient and locally-persistent species across all sites. For any traits with significant overall differences, we performed linear regressions to see whether the magnitude of trait-based differences between locally-transient and locally-persistent species trended with site temperature or precipitation. We did not test for trait-based differences by species status in seedling communities because the number of locally-transient species was too low to provide any confidence in calculations of within-site trait means (only 4 ± 3 locally-transient species occurred on average as emerged seedlings at each site; Appendix S1: Table S1). All scripts in this study were written in R and are available at https://github.com/guittarj/MS_Transients.

## RESULTS

### Mature vegetation rarefactions

Rarefaction analysis indicated that the numbers of plots of mature vegetation surveyed at each site were more than sufficient to stabilize the local composition of locally-persistent species (Appendix S1: Fig. S2), lending confidence to our locally-transient/locally-persistent species status assignments. Furthermore, no locally-transient species by itself ever represented more than 0.4% of total cover at any site (Appendix S1: Fig. S3), illustrating the minor overall contribution of locally-transient species to local community structure.

### Evidence for local filtering

Seeds of locally-transient species occurred at all 12 of our grassland sites, representing, on average, 4 of 42 species in the combined seed rain and seed bank. In the combined seed rain and seed bank, the number of locally-transient species and their total abundances increased significantly with temperature (Fig. 2; species richness: p = 0.045, *R^2^* = 0.34; total abundance: p = 0.016, *R^2^*= 0.46). These trends were driven primarily by increases of local-transients in the seed bank (species richness: p = 0.031, *R^2^*= 0.39; total abundance p = 0.035, *R^2^* = 0.37), not the seed rain (Appendix S1: Fig. S4), underscoring the fact that the composition of the seed rain likely differs between years, and that the seed bank likely serves as an important reservoir of local plant diversity. There were no significant trends in species richness or total abundance with precipitation in either the seed rain, the seed bank, or the two combined. Species richness in the adult vegetation rose significantly with temperature (p = 0.0063, *R^2^* = 0.54) but not with precipitation (p = 0.871).

**Figure 2.**
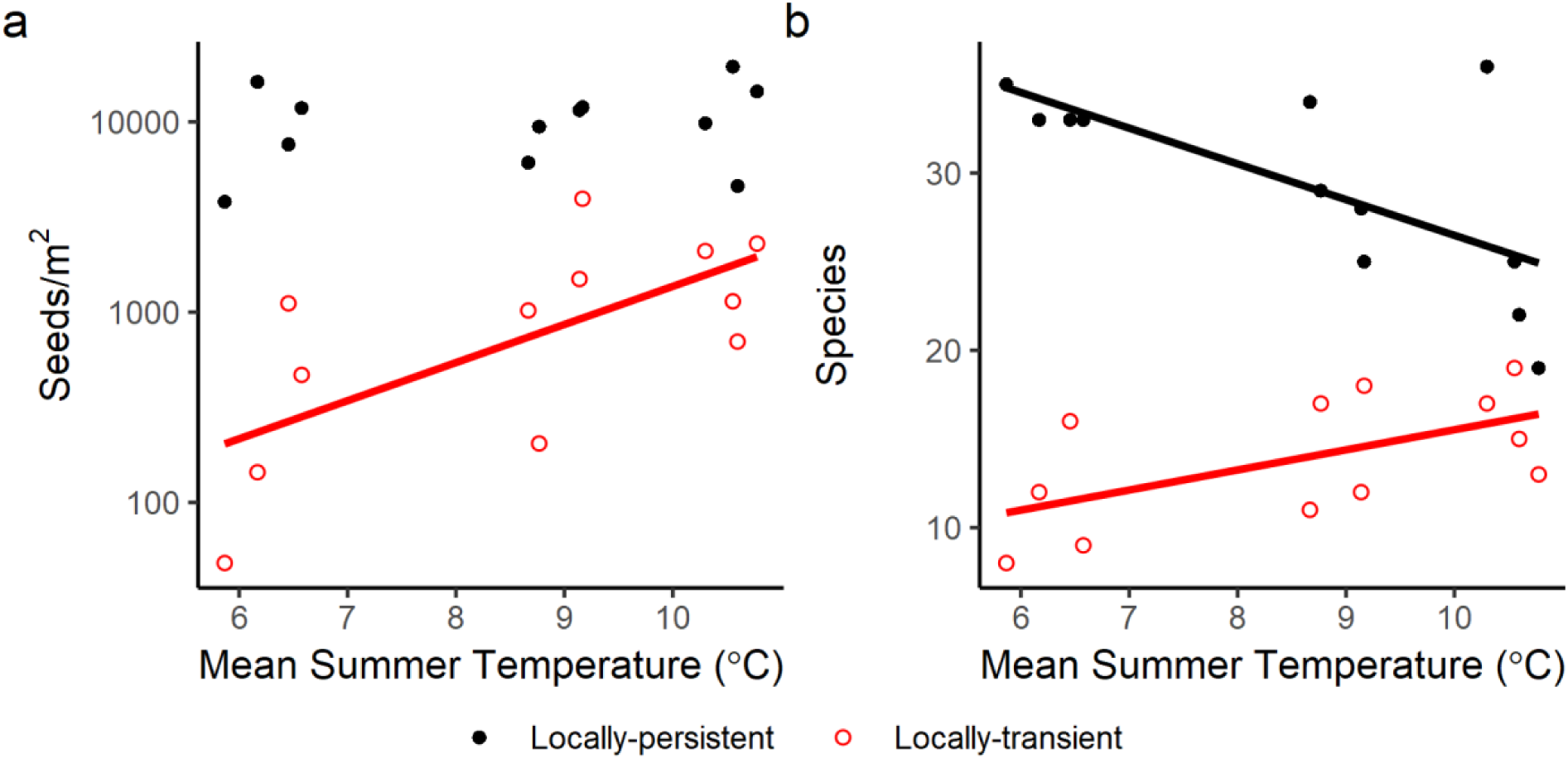
Seed density (a) and species richness (b) in the combined seed rain and seed bank, grouped by locally-transient/locally-persistent species status and plotted by mean summer temperature. Summer is defined as the four warmest months at each site. Solid lines are present when temperature was a significant predictor (p < 0.05) of log_10_-transformed seed density or species richness in a linear regression. A parallel set of regressions was performed using mean annual precipitation as the sole predictor, but no trends were even marginally significant (p-values always > 0.1).

About 10% of all seeds were from locally-transient species, with more transients represented in the seed bank (ca. 13% of total seed bank) than in the seed rain (ca. 4% of total seed rain) (Appendix S1: Table S2). While most locally-transient populations had locally-persistent adult populations at sites with similar climates, some did not (Appendix S1: Table S2). Operating on the assumption that locally-transient species dispersed from the most-climatically similar sites at which they persist as adults, seeds of locally-transient species were about four times more likely to have dispersed outside of their realized climate niches into warmer sites (i.e., from cooler climates) and slightly more likely to have dispersed outside of their realized climate niches into drier sites (i.e., from wetter climates) (Appendix S1: Table S2).

To determine how sensitive our conclusions were to changes in our operational definition of locally-transient/locally-persistent species status, we explored how results changed under each of the four possible cutoff scenarios offered by our data. Specifically, we looked at how results changed as the definition of locally-persistent species shifted from those species present in at least one, at least two, at least three, or all four annual surveys of mature vegetation at each given site. As the cutoff for locally-persistent became more stringent, and the cutoff for locally-transient (by definition) relaxed, the total number of locally-transient populations (i.e., unique species*site combinations) in the combined seed rain and seed bank at our sites rose from 119 (1989 seeds), to 149 (2549 seeds), to 167 (3665 seeds), to 205 (5007 seeds). However, these differences in the estimated numbers of locally-transient/locally-persistent species by site did not alter our main conclusions that (1) the influence of local filtering increases with temperature, and that (2) grassland sites in southern Norway are connected by dispersal, albeit primarily among nearby sites with similar climates.

### The stage-wise removal of locally-transient species

Locally-transient species were outperformed by locally-persistent species during seedling emergence (Fig. 3, Table 1), but not seedling establishment (Fig. 3, Appendix S1: Table S3). The lower emergence rates of locally-transient species appeared to be driven primarily by species which had putatively dispersed outside of their realized climate niches into cooler climates, drier climates, or those that had dispersed from unknown climates (see “Origin-based predictors” in Table 1). In addition, after accounting for local seed abundances, warmer sites tended to have higher rates of seedling emergence than cooler sites (“General predictors” in Table 1), linking climate and overall seedling performance. No GLM of seedling establishment outperformed null expectations (Appendix S1: Table S3), suggesting that neither climate, species status, nor putative seed origins were meaningful predictors of seedling establishment rates.

**Figure 3.**
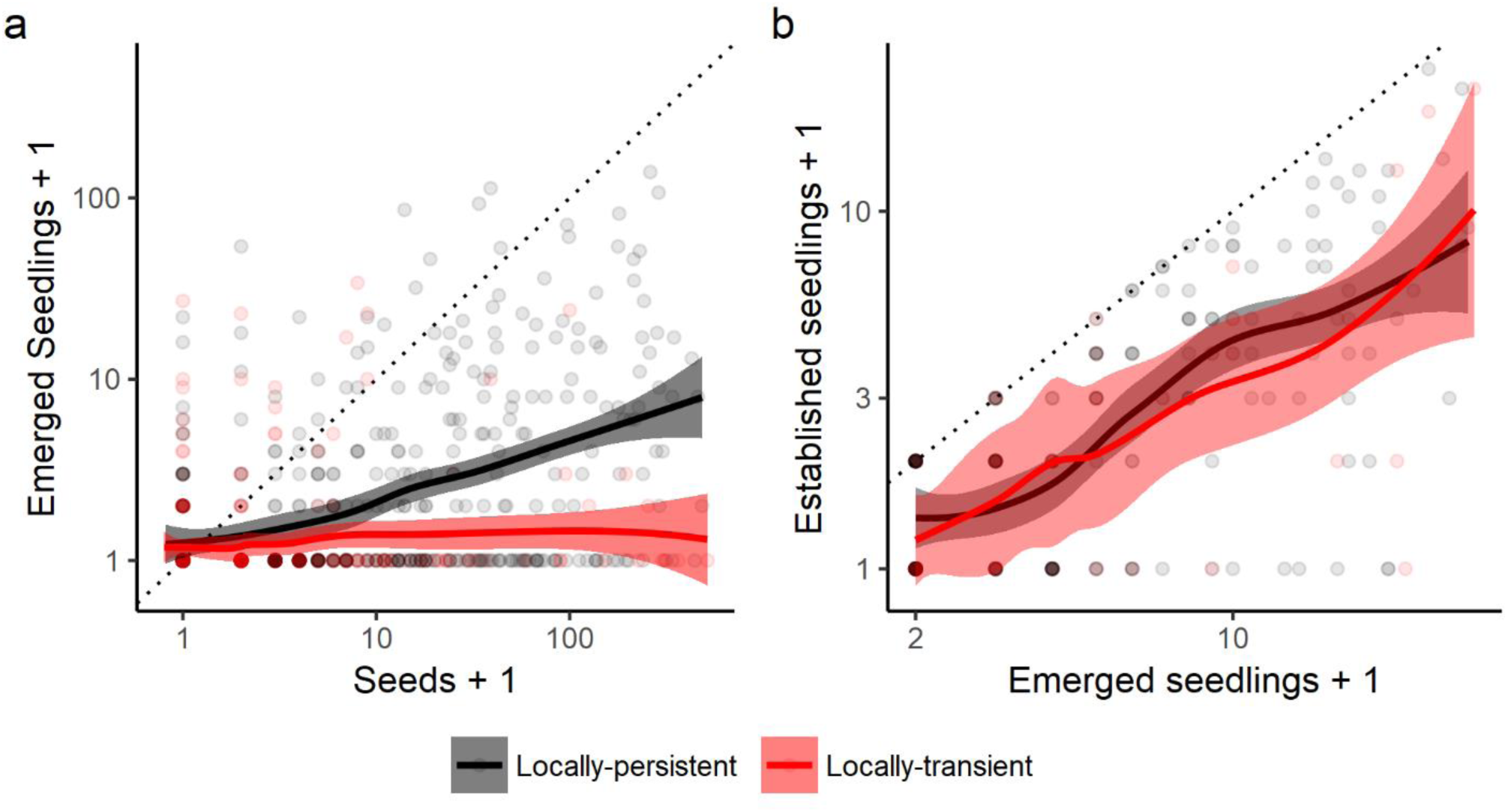
Life stage transition probabilities, grouped by locally-transient/locally-persistent species status. Specifically, numbers of emerged seedlings of each species at each site are plotted by their corresponding number of seeds at each site (a), and numbers of established seedlings of each species at each site are plotted by their corresponding number of emerged seedlings at each site (b). As such, each circle represents the presence/absence of one species at one site, and colored lines and shadings show LOESS smoothing functions and 95 % confidence intervals. Seed abundances equal the total number of seeds in the seed rain and seed bank. Seedling abundances equal the total number of individuals in four 25 x 25 cm subplots at each site. Count data are increased by one to allow for plotting zeroes on a log scale. Panels only show data falling within the observed window of locally-transient seed abundances (< 400 seeds) and locally-transient seedling abundances (< 33 seedlings) in order to focus on the comparison of locally-transient and locally-persistent species.

**Table 1.**
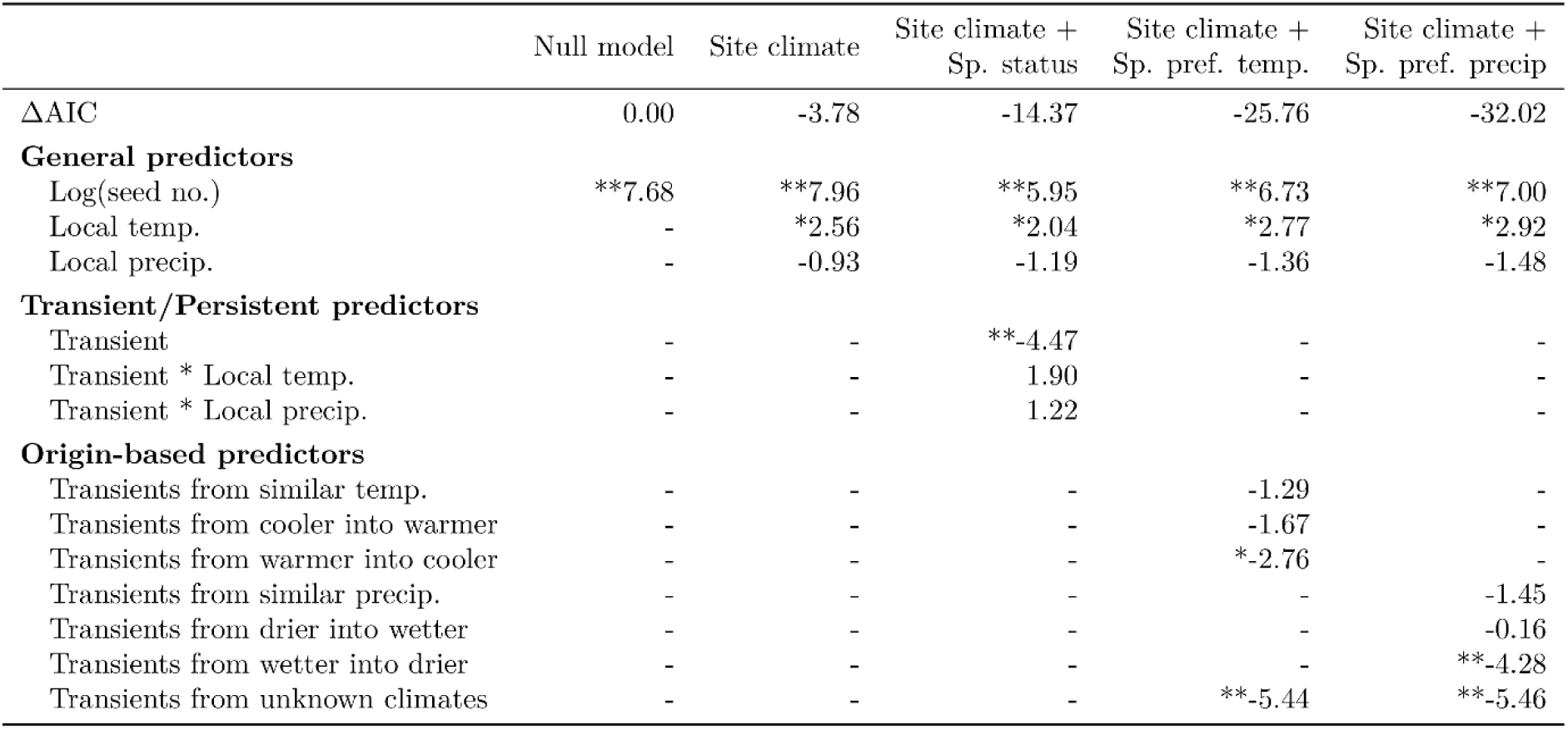
Standardized coefficients (z-scores) from different GLM models (columns) predicting numbers of emerged seedlings by species and site. In column headers (i.e., model descriptions), “Species status” refers to whether the species is locally-transient or locally-persistent, and “Species preferred temp/precip” refers to the nearest climates at which the species has a persistent adult population, which we use to infer the climate from which they likely dispersed. Data include all recorded seeds and seedlings that could be identified to species. N is equal to the number of unique species-by-site combinations for each category/predictor. Asterisks denote significance (*: p < 0.05, **: p < 0.001). Dashes indicate predictors that were not included in a given model. Including model terms for locally-transient/locally-persistent species status and then preferred temperature and precipitation progressively improved model performance, as reflected in the reduction of AIC values relative to the null model that uses only local seed numbers as a predictor.

To confirm that these results were not artifacts of how we combined seed rain and seed bank data (e.g., if seeds of locally-transient species emerged at lower rates because most were from the seed bank, and the seed bank had overall lower rates of emergence), we re-ran GLMs with only seed rain data and observed qualitatively similar results (data not shown). To determine how sensitive our conclusions involving seedling performance were to changes in the operational definition of locally-transient/locally-persistent species status, we re-ran the GLMs for emergence and establishment using each of the four possible locally-transient/locally-persistent cutoff scenarios offered by our data. Changes to the locally-transient/locally-persistent cutoff did not alter our overall conclusion that locally-transient species are disfavored during emergence but not establishment (Appendix S1: Fig. S5).

### Trait-based mechanisms of local filtering

Locally-transient and locally-persistent species in the combined seed rain and seed bank differed significantly in three functional traits (Fig. 4). Specifically, when averaged across species at the site level, locally-transient species were significantly shorter, had smaller seeds, and had longer-lasting vegetative connections among ramets than locally-persistent species. Of the three traits that differed consistently between locally-transient and locally-persistent species, the only instance where those differences varied significantly with site climate was an increase in the magnitude by which locally-transient species had longer-lasting connections than locally-persistent species with increasing site temperature.

**Figure 4.**
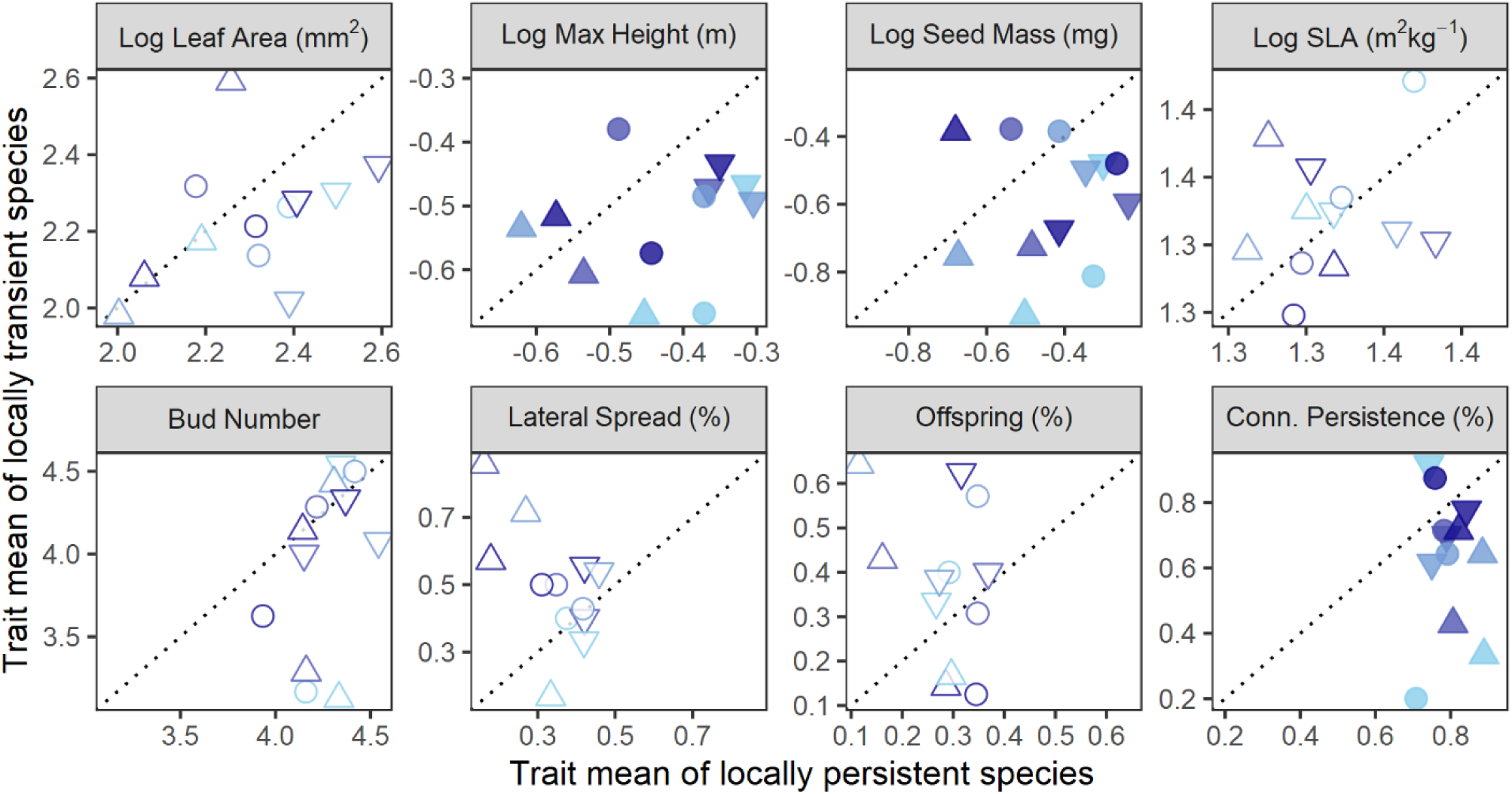
Mean trait values of locally-transient species relative to locally-persistent species at each site. Panels show data for four commonly used traits (top row) and four clonal traits (bottom row). Shapes and shadings are consistent with Fig. 1 and reflect approximate mean summer temperatures of 6 °C (triangle), 9 °C (circle), and 10.5 °C (inverted triangle) and annual precipitations of 650 mm, 1300 mm, 2000 mm, and 2900 mm, from light blue to dark blue. Shapes are filled with color only when values differ significantly between locally-transient and locally-persistent species across all sites (paired t-test; p < 0.05). SLA: specific leaf area. Bud number is an integer score that ranges from 1 (few buds either belowground or aboveground) to 8 (many buds both below and aboveground); percent lateral spread refers to the proportion of species with rates of lateral spread greater than 1 cm/year; percent offspring refers to the proportion of species that commonly produce two or more vegetative offshoots per parent per year; percent connection persistence refers to the proportion of species with inter-ramet connections that persist for two or more years.

## DISCUSSION

In this study, we compared the compositions of seed, seedling, and adult plant communities along temperature and precipitation gradients to shed light on how seed dispersal and local filtering processes interactively generate and maintain patterns of community composition at the landscape scale. Overall, our results point to four main conclusions: (1) the influence of local filtering on community membership increases with temperature, (2) local filtering is stronger during seedling emergence than during seedling establishment, (3) climate-based niche differences drive differential performance among species, especially for seeds dispersing outside of their realized climate niches into colder and drier climates (i.e., dispersing from warmer and wetter climates), and (4) locally-transient species have traits which may make them better dispersers (smaller seeds) but poorer competitors for light (shorter statures, less persistent clonal connections), which helps explain their arrival to new sites but subsequent failure to establish persistent adult populations. We elaborate on these conclusions below, and end with a discussion of what our results mean for how these alpine grasslands will likely respond to climate change.

### Community assembly and the determinants of species ranges

Community membership and the potential for compositional change over time are significantly influenced by the set of seeds dispersing, or not dispersing, to a given site. We found relatively few species to be dispersing outside of their realized climate niches, highlighting the potentially important role that dispersal limitation plays in determining community composition, at least on shorter time scales. At the same time, the few species that successfully dispersed outside of their realized climate niches were the ones most strongly disfavored by local filtering processes (Table 1), highlighting the important role that local filtering also plays in determining community composition in our system. In other words, our results explicitly show how dispersal dynamics and local filtering interactively influence community assembly.

Within the overall result of local filtering (Table 1), we observed a significant increase in the fraction of the seed community that was locally-transient from the coldest (highest altitude) to warmest (lowest altitude) sites (Fig. 2), suggesting that the influence of local filtering on community membership increases with temperature. Such a trend with temperature could arise for at least three non-exclusive reasons. First, warm-adapted species could have spatially larger seed dispersal shadows than cold-adapted species, resulting in more seeds traveling to unsuitable patches, such as those with undesirable topographic, edaphic, or biotic conditions. Factors affecting seed dispersal distances include wind speed and plant height (Thomson et al. 2011), although neither are likely to be important in our experimental system: the colder, alpine sites are consistently windier than the warmer, lowland sites, due to their more exposed topography and less protective tree-cover, and the variation in mean maximum height among our sites is minor (< 30 cm) (Guittar et al. 2016) compared to the differences in plant height known to affect dispersal distance (Thomson et al. 2011). A second explanation for the putative increase in the strength of filtering with temperature could be that seeds are simply more likely to disperse downhill (i.e., from colder high-altitude sites to warmer low-altitude sites) than the reverse, again leading to more seeds arriving at unsuitably warm patches than unsuitably cold patches. A third explanation could be that competition for resources is fiercer at warmer temperatures, resulting in more local extirpations in the adult plant communities at warmer sites, although, importantly, this explanation presumes that similar numbers of seeds and species are dispersing to all sites, which may not be the case. Nonetheless, there is support such an explanation: competition for light is hypothesized to be stronger at warmer sites due to higher productivity, more developed soils, and overall less stressful growth conditions (Grime 1973, Olsen et al. 2016). In our study, species richness fell steeply with temperature in the adult vegetation (from an average of 42 species at the warmest sites to 64 species at the coldest sites), but only slightly in the seed rain and not in the seed bank (Fig. 2), consistent with a scenario in which similar numbers of species arrived to all sites, but more species were competitively excluded by adulthood at warmer sites.

Surprisingly, locally-transient species were significantly disfavored during seedling emergence (Fig. 3, Table 1), but not during seedling establishment (Fig. 3, Appendix S1: Table S3), suggesting that germination and very early survival is a more important determinant of seedling success than successful establishment in our system. This result contradicts conventional expectations of weak filtering during seedling emergence but strong filtering during seedling establishment (Moles and Westoby 2004). Differential germination rates among species could be driven by species preferences for soil (Evans and Etherington 1990, Benvenuti 2003) or climate conditions. It should be noted that while some individuals could have been filtered out after meeting our operational definition of establishment (i.e., had stems at least 2 cm long and, if a forb, also had non-cotyledonous leaves), but before becoming reproductively active (i.e., before establishment *sensu stricto*), this does not appear to be the case. Across sites, the mean proportion of species that were locally-transient among established seedlings was only slightly higher than that found in the adult vegetation data (∼14.6 % vs. ∼13.6 %, respectively), indicating that any locally-transient species which successfully emerged were essentially just as likely as locally-persistent species to survive long enough to be captured by our mature vegetation surveys, but not long enough to be considered locally-persistent. In other words, locally-transient species evidently failed to persist as adults not because they failed to establish, but either because they failed to germinate or because they failed to compete as post-establishment juveniles, resulting in local population growth rates below maintenance levels.

The low emergence rates among locally-transient species were largely driven by seeds that had putatively dispersed outside their realized climate niches into cooler climates (i.e., from warmer climates), into drier climates (i.e., from wetter climates), and those from unknown climates (Table 1), illustrating how climate-based niche preferences can influence community membership and thus range expansion. Climate is known to play a pivotal role in the release of seed dormancy in plants (Probert 2000), and variation in germination timing is known to occur even among populations of the same species at different climates (Shimono and Kudo 2003, Bischoff et al. 2006, Spindelböck et al. 2013). If the seeds of locally-transient species dispersing outside of their realized climate niches into colder and/or drier climates were predisposed to emerging before spring is safely underway, this could explain their particularly low emergence rates. A second, complementary explanation could be that seeds dispersing outside of their realized climate niches into colder and/or drier climates simply find these conditions more stressful and are thus less likely (i.e., not adapted) to successfully emerge as seedlings.

Locally-transient species differed consistently from locally-persistent species in their functional traits, offering some mechanistic hypotheses for why they can disperse into new sites but fail to persist as adults (Fig. 4). First, locally-transient species had consistently smaller seeds than locally-persistent species, which is thought to increase dispersal distance (Greene and Johnson 1993, Westoby 1998), and may also decrease performance upon arrival due to their smaller maternal subsidies (Moles and Westoby 2006). Second, locally-transient species were shorter and had less long-lasting ramet-ramet connections, on average, than locally-persistent species, suggesting that the ability to grow both vertically and exchange resources horizontally confer important advantages for long-term survival in mature vegetation. Maximum potential height and the capacity for clonal growth are associated with the ability to compete for light (Falster and Westoby 2003) and soil resources in spatially heterogenous systems (Oborny and Kun 2003, Eilts et al. 2011), respectively, and can work synergistically to confer competitive ability in herbaceous communities (Gough et al. 2012). These two traits were significantly associated with competitive ability in a turf transplant experiment in the same system (Guittar et al. 2016), offering additional support for the importance of architectural traits in driving grassland species performance. Equally noteworthy were the generally weak correlations between locally-transient and locally-persistent species in the remaining functional traits. The weak correlations in these traits does not imply that they have no influence on the selective removal of locally-transient species, but that their potential influence is not consistent or predictable across sites in the region.

The trait-based differences between locally-transient species and locally-persistent species align with a general tradeoff between colonization and competitive ability (Levins and Culver 1971, Tilman 1994, Amarasekare and Nisbet 2001, Yu and Wilson 2001), and thus suggest that such a tradeoff, when combined with sufficient levels of disturbance, is an important driver of succession and community assembly dynamics. That is, the smaller seeds of locally-transient species may promote their probability of arriving to fresh openings in the canopy, while their lower statures and decreased capacity for clonal growth may decrease their chances of establishing locally-persistent populations. While our trait-based analysis offers no obvious mechanistic explanation for why locally-transient species were disfavored during seedling emergence, the trait-based differences may correlate with unmeasured traits that do influence species performance during seedling emergence, and thus may serve as important proxies.

### Implications for community responses to climate and climate change

Community response to climate change will depend both on species’ abilities to track environmental changes via dispersal, and on niche-based performance differences among species in different environments (Graae et al. 2017). Southern Norway is expected to grow warmer and wetter as climate change progresses (Hanssen-Bauer et al. 2009), so species will have to disperse into cooler (upslope) and drier (more inland) sites to maintain their current climate conditions. However, of the 122 species with persistent adult populations at one or more of our sites, only 10 species (representing 0.3 % of total seeds) dispersed outside of their realized climate niches into cooler sites, and 13 species (representing 1.9 % of total seeds) into drier sites (Appendix S1: Fig. S4, Appendix S1: Table S2), suggesting that many if not most species will likely be unable to rapidly shift their populations into cooler and drier climates, and are at risk of being extirpated as competitively-superior species arrive to their communities (Alexander et al. 2015). Yet, our finding that immigrant seeds from other climates were strongly disfavored provides a potential silver lining: if a species can disperse into a site with a newly suitable climate, it should be strongly favored over local residents, and thus be able to quickly establish a locally-persistent population. Overall, our results underscore the dynamic way in which communities will likely respond to climate change, and emphasize the need for further work on how species will vary in their ability to disperse and compete in different community contexts.

## Acknowledgements

We thank the landowners for allowing us to set up the experiments on their land, the Norwegian Research Council (KLIMAFORSK project 184912/S30), NSF GRFP (JLG; Grant Number: 1256260), and Norge-Amerika Foreningen for funding. We thank Siri L. Olsen, Mari Jokerud, Pascale Michel, Hilary H. Birks and Berhe Luel for contributions to the data collection.

## Authors’ contributions

JG wrote the manuscript and analyzed data with conceptual and editorial help from all authors. VV, KK, and DG conceived of the experimental study system and the experiments upon which this paper is based. VV, KK, and AB gathered seedling and mature vegetation data, MB gathered seed rain data, and KK, EM, JT, AB, and VV gathered seed bank data. VV and JG secured funding.

