## Supplemental figures and tables for "Dispersal dynamics and local filtering vary with climate across a grassland landscape" (from another version of this preprint: version 1, 2019-03-05)

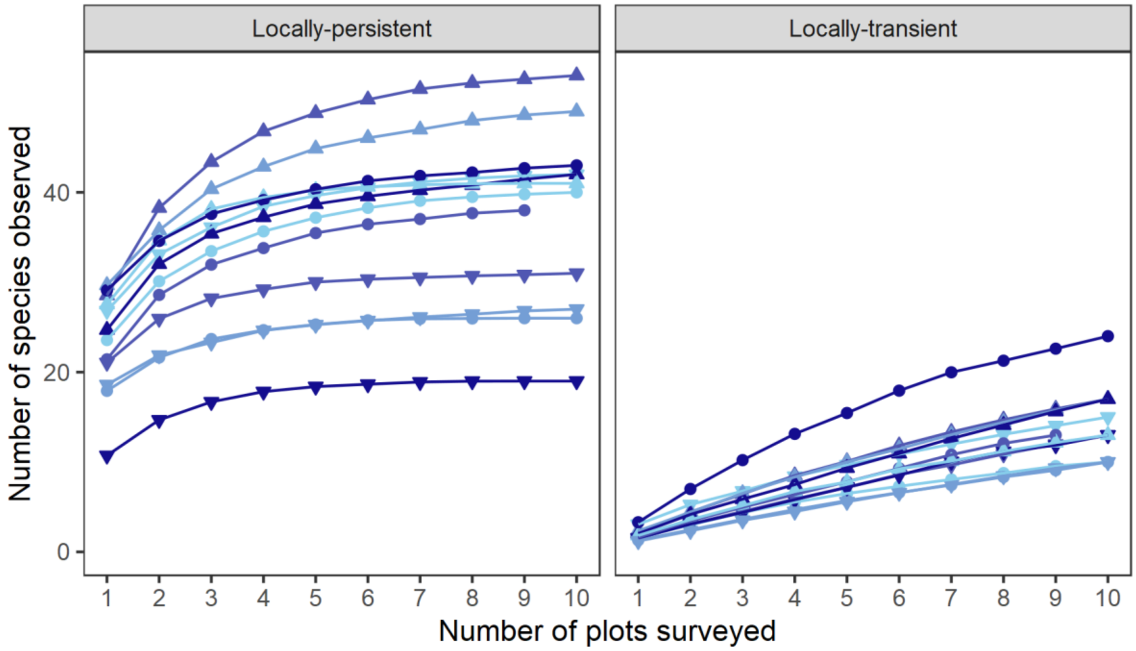


**Figure S1.** Rarefaction curves show increases in the number of species observed in the mature vegetation at each site with increasing sampling effort, grouped by local transient/persistent species status. Species occurrence data for each 25 x 25 cm plot were determined by pooling data from four annual vegetation surveys conducted at peak biomass at each site in 2009, 2011, 2012, and 2013. Species were considered locally-persistent if observed in any plot at a site in at least three of the four vegetation surveys, otherwise species were considered locally-transient. Rarefaction was performed 100 times, with the stepwise order of plots shuffled randomly each time. Colored shapes show mean values across the 100 replicates. Each line is a site. Shapes and shadings are consistent with Fig. 1 and reflect approximate mean summer temperatures of 6 °C (triangle), 9 °C (circle), and 10.5 °C (inverted triangle) and annual precipitations` of 650 mm, 1300 mm, 2000 mm, and 2900 mm, from light blue to dark blue.

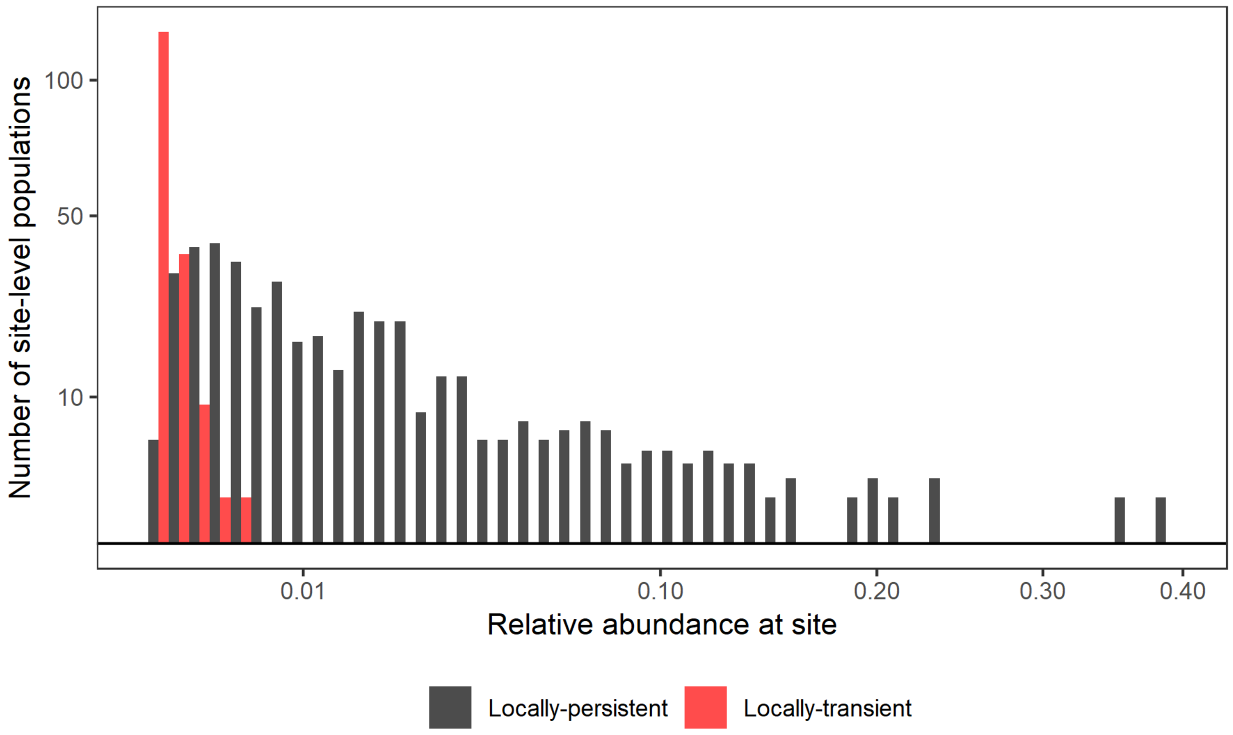


**Figure S2.** A histogram of local relative abundances of all species at each of all twelve sites, grouped and colored by local transient/persistent species status, and plotted along square-root scales on both x and y axes. All locally-transient species fall below 0.31% total site cover.

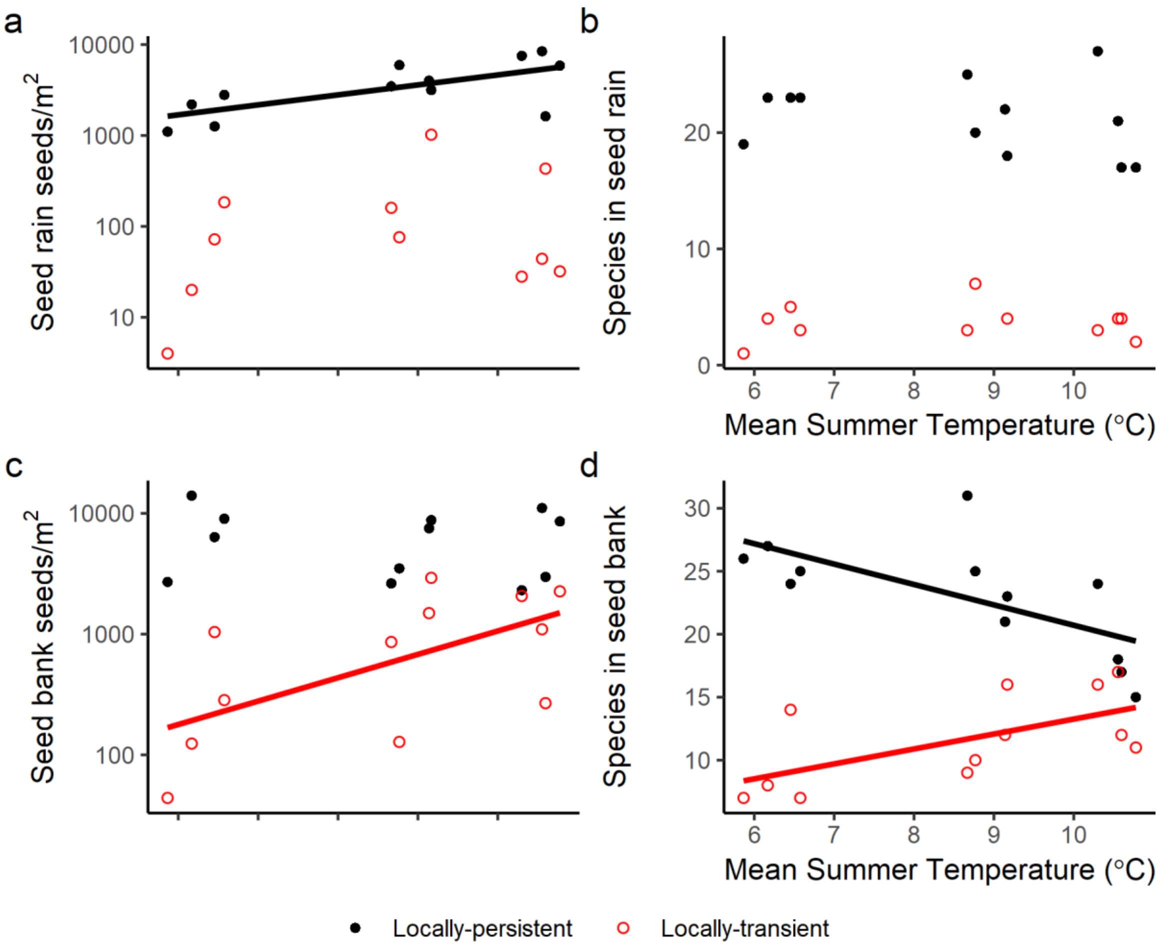


**Figure S3.** Seed density in the seed rain (a) and seed bank (c), and species richness in the seed rain (b) and seed bank (d), grouped by locally-transient/locally-persistent species status and plotted by mean summer temperature. Summer is defined as the four warmest months at each site. Solid lines are present when temperature was a significant predictor ($p < 0.05$) of $\log_{10}$-transformed seed density or species richness in a simple linear regression. A parallel set of regressions was performed using mean annual precipitation as the sole predictor, but no trends were even marginally significant (p-value was always > 0.1).

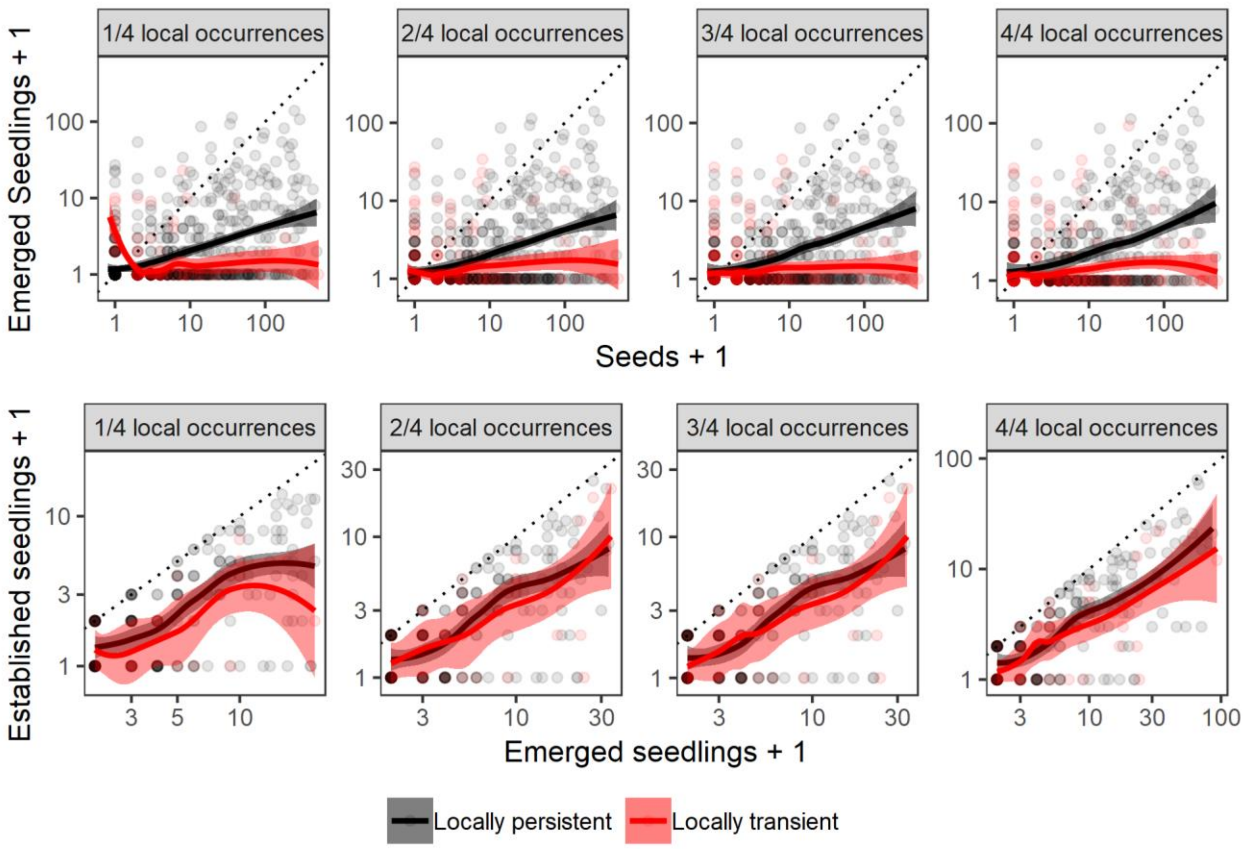


**Figure S4.** The abundances of emerged seedlings (top row of panels) and established seedlings (bottom row of panels) of species at individual sites, plotted by local seed number and local seedling number, respectively, and colored by local species status. Each column of panels reflects a version of the data for which a different cutoff for local persistent species status was used; from left to right, only species that occurred in the mature vegetation at a given site in one, two, three, or four of the four total annual surveys were considered locally "persistent," and the remaining species (under each cutoff scenario) were considered locally "transient." Each circle represents the presence/absence of one species at one site, and colored lines and shadings show LOESS smoothing functions and 95 % confidence intervals. Seed number is equal to the sum of seeds in the seed rain and seed bank. Abundances reflect the total number of individuals in four 25 x 25 cm subplots at each site. Count data are increased by one to include zeroes on a log scale. Panels only show data falling within the observed window of locally-transient seed abundances and locally-transient seedling abundances under each cutoff scenario in order to focus on the comparison of locally-transient and locally-persistent species. These figures illustrate that regardless of the transient/persistent cutoff used, (1) emergence rates remained significantly lower for seeds of transient species than seeds of persistent species (top row of panels), and (2) establishment rates did not significantly differ among seedlings of transients and persistent species (bottom row of panels). Model selection and coefficient significance values were also effectively the same among the four scenarios (data not shown).

| Stage | Density (per sq. m.) | Site richness | Regional richness |
|---|---|---|---|
| **All individuals** | | | |
| Seed Rain | 4189 ± 2456 | 25 ± 4 | 94 |
| Seed Bank | 7715 ± 4045 | 35 ± 4 | 103 |
| All Seeds | 11905 ± 5041 | 43 ± 5 | 126 |
| Emerged Seedlings | 1459 ± 613 | 20 ± 5 | 82 |
| Established Seedlings | 415 ± 206 | 14 ± 4 | 65 |
| Seeds and Seedlings | 13778 ± 5567 | 46 ± 5 | 129 |
| Adults | | 53 ± 13 | 144 |
| All Stages | | 63 ± 11 | 158 |
| **Individuals of locally-transient species only** | | | |
| Seed Rain | 176 ± 296 | 3 ± 2 | - |
| Seed Bank | 1053 ± 970 | 12 ± 4 | - |
| All Seeds | 1229 ± 1132 | 14 ± 4 | - |
| Emerged Seedlings | 331 ± 418 | 4 ± 3 | - |
| Established Seedlings | 61 ± 94 | 2 ± 2 | - |
| Seeds and Seedlings | 1621 ± 1484 | 16 ± 5 | - |

**Table S1.**

Densities and species richness values within and across sites for each life stage. The density of ”all individuals”” includes unidentified seeds and seedlings. Regional richness is not shown for locally-transient species because locally-transient/locally-persistent species status can vary by site, and thus cannot be summarized in this way across sites.

| Species status | Seed rain | | Seed bank | | All seeds | | Emerged | | Established | |
|---|---|---|---|---|---|---|---|---|---|---|
| | **No.** | **%** | **No.** | **%** | **No.** | **%** | **No.** | **%** | **No.** | **%** |
| All individuals | 12370 | 100.0 | 23014 | 100.0 | 35384 | 100.0 | 3039 | 100.0 | 1079 | 100.0 |
| Locally-persistent | 11852 | 95.8 | 19867 | 86.3 | 31719 | 89.6 | 2778 | 91.4 | 983 | 91.1 |
| Locally-transient | 518 | 4.2 | 3147 | 13.7 | 3665 | 10.4 | 261 | 8.6 | 96 | 8.9 |
| **Temperature of nearest persistent adult population** | | | | | | | | | | |
| Same | 428 | 3.5 | 2061 | 9.0 | 2489 | 7.0 | 180 | 5.9 | 83 | 7.7 |
| Cooler | 16 | 0.1 | 356 | 1.5 | 372 | 1.1 | 41 | 1.3 | 2 | 0.2 |
| Warmer | 21 | 0.2 | 72 | 0.3 | 93 | 0.3 | 5 | 0.2 | 5 | 0.5 |
| Unknown | 53 | 0.4 | 658 | 2.9 | 711 | 2.0 | 35 | 1.2 | 6 | 0.6 |
| **Precipitation of nearest persistent adult population** | | | | | | | | | | |
| Same | 154 | 1.2 | 1636 | 7.1 | 1790 | 5.1 | 155 | 5.1 | 58 | 5.4 |
| Drier | 51 | 0.4 | 441 | 1.9 | 492 | 1.4 | 62 | 2.0 | 28 | 2.6 |
| Wetter | 260 | 2.1 | 412 | 1.8 | 672 | 1.9 | 9 | 0.3 | 4 | 0.4 |
| Unknown | 53 | 0.4 | 658 | 2.9 | 711 | 2.0 | 35 | 1.2 | 6 | 0.6 |

**Table S2.**
Numbers of individuals recorded, grouped by locally-transient/locally-persistent species status (top three rows). Then, locally-transient individuals are further grouped by the putative climate preferences of their species, i.e., the temperatures/precipitations of the nearest sites at which persistent adult populations are known to occur, relative the climates of the local sites. Percentages are of all individuals (top row). Individuals that could not be identified to species were not included.

| | Null model | Site climate | Site climate + Sp. status | Site climate + Sp. pref. temp. | Site climate + Sp. pref. precip |
|---|---|---|---|---|---|
| ΔAIC | 0.00 | 3.59 | 0.05 | 0.29 | 10.27 |
| **General predictors** | | | | | |
| Log(seedling no.) | **18.92 | **18.76 | **18.48 | **18.49 | **18.20 |
| Local temp. | - | 0.51 | 1.36 | 1.07 | 0.64 |
| Local precip. | - | -0.46 | 0.05 | -0.19 | -0.60 |
| **Transient/Persistent predictors** | | | | | |
| Transient | - | - | 0.74 | - | - |
| Transient * Local temp. | - | - | *-2.97 | - | - |
| Transient * Local precip. | - | - | -1.48 | - | - |
| **Origin-based predictors** | | | | | |
| Transients from similar temp. | - | - | - | -0.04 | - |
| Transients from cooler into warmer | - | - | - | *-2.63 | - |
| Transients from warmer into cooler | - | - | - | 1.39 | - |
| Transients from similar precip. | - | - | - | - | -0.84 |
| Transients from drier into wetter | - | - | - | - | 0.33 |
| Transients from wetter into drier | - | - | - | - | 0.18 |
| Transients from unknown climates | - | - | - | -0.73 | -0.70 |

**Table S3.**
Standardized coefficients (i.e., z-scores) from different GLM models (columns) predicting numbers of established seedlings by species and site. In column headers (i.e., model descriptions), "Species status" refers to whether the species is locally-transient or locally-persistent, and "Species preferred temp/precip" refers to the nearest climates at which the species has a persistent adult population, which we use to infer the climate from which they likely dispersed. Data include all recorded seedlings that could be identified to species. Asterisks denote significance (*: $p < 0.05$, **: $p < 0.001$). Dashes indicate predictors that were not included in a given model. Model terms for locally-transient/locally-persistent species status and preferred temperature or precipitation did not improve model performance, as reflected in the lack of any decrease in AIC values relative to the null model based only on local numbers of emerged seedlings.
